# Temporal analysis of localization and trafficking of glycolipids

**DOI:** 10.1101/2020.04.07.030411

**Authors:** Kenta Arai, Kazuya Kabayama, Yoshimi Kanie, Osamu Kanie, Koichi Fukase

## Abstract

Glycolipid metabolism occurs in the Golgi apparatus, but the detailed mechanisms have not yet been elucidated. We used fluorescently labeled glycolipids to analyze glycolipid composition and localization changes and shed light on glycolipid metabolism. In a previous study, the fatty chain of lactosyl ceramide was fluorescently labeled with BODIPY (LacCer-BODIPY) before being introduced into cultured cells to analyze the cell membrane glycolipid recycling process. However, imaging analysis of glycolipid recycling is complicated because of limited spatial resolution. Therefore, we examined the microscopic conditions that allow the temporal analysis of LacCer-BODIPY trafficking and localization. We observed that the glycolipid fluorescent probe migrated from the cell membrane to intracellular organelles before returning to the cell membrane. We used confocal microscopy to observe co-localization of the glycolipid probe with endosomes and Golgi markers, demonstrating that it recycles mainly through the trans-Golgi network (TGN). Here, a glycolipid recycling pathway was observed that did not require the lipids to pass through the lysosome.

## Introduction

The homeostasis of glycolipids expressed on the cell membrane is maintained by a balance between de novo synthesis and degradation and recycling (1-3). Several studies have biochemically analyzed these pathways using inhibitors (4, 5). Pagano *et al*. performed a spatio-temporal analysis of glycolipid metabolism by adding fluorescently labeled glycolipids to cells and fractionating organelles. They used fluorescently labeled lactosyl ceramide (LacCer-BODIPY, Fig. 1A) in which the fatty acid chain was replaced with a 5-carbon alkyl chain linked to the fluorescent probe BODIPY. They confirmed that this probe was taken up by cells via the same route as radiolabeled LacCer (6-12). These reports demonstrated that glycolipids internalized from the cell membrane localize to the Golgi apparatus and lysosomes before some glycolipids are recycled back to the cell membrane. However, a comprehensive spatial and temporal understanding of the transported glycolipids has not yet been achieved due to differences in the conditions and analytical methods of adding these glycolipid derivatives.

**Figure 1.**
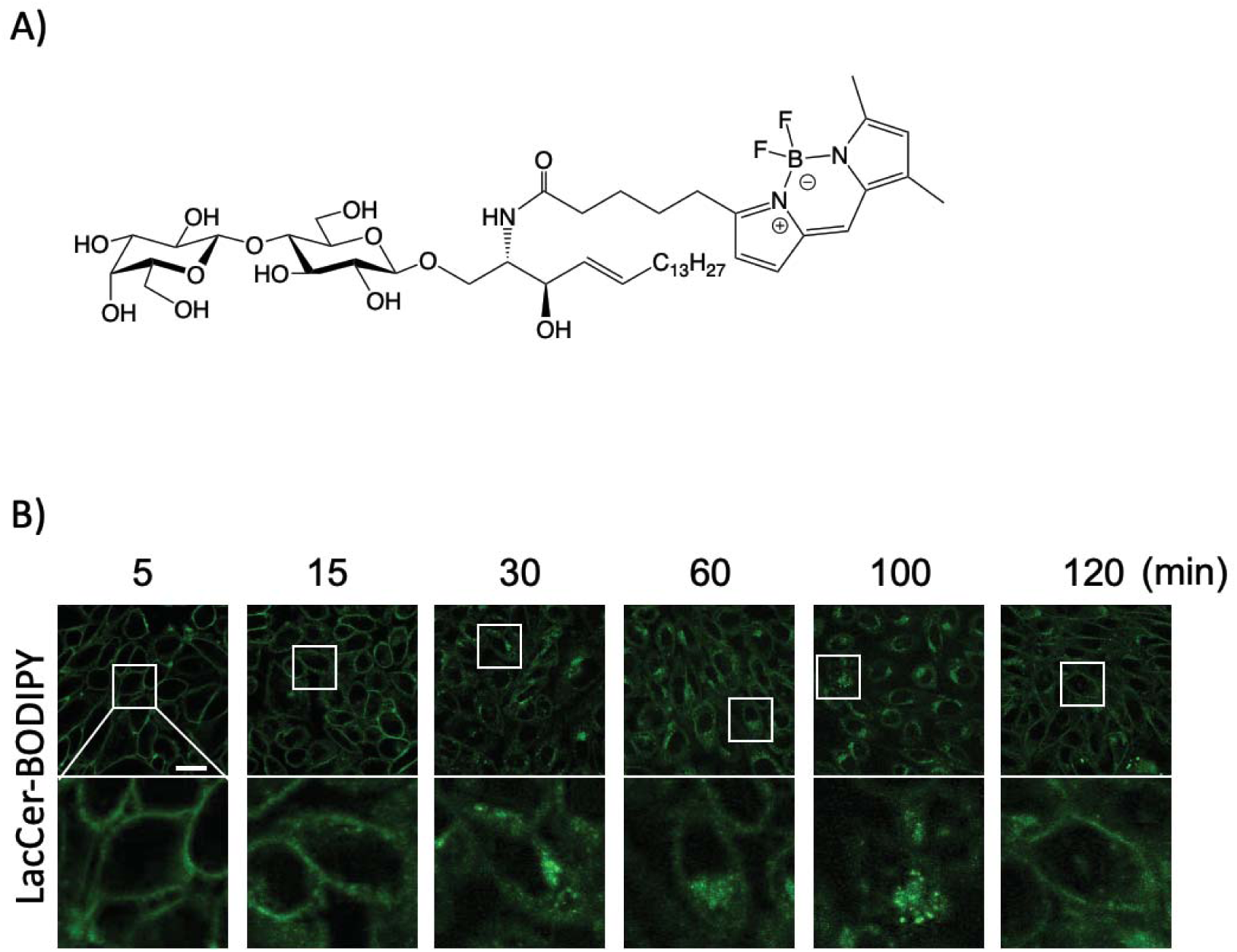
Time-dependent localization of LacCer-BODIPY using pulse chase in CHO-K1 cells. A) Structure of BODIPY C_5_ labeled lactosyl ceramide (LacCer-BODIPY). B) Confocal microscope images of LacCer-BODIPY localization when incubated at 5, 15, 30, 60, 100, and 120 min after 1-min pulse labeling. The lower panels show enlargement of the area within the white-box of the upper panels. Scale bar; 10 µm.

We previously constructed a system to spatio-temporally analyze glycolipid metabolism using LacCer-BODIPY (13-15). We used this exogenous glycolipid probe to perform a detailed time-lapse analysis of the recycling pathway. Here, we successfully visualized a transport pathway that translocates glycolipids to the Golgi without them having to pass through the lysosome and whereby they quickly return to the cell membrane.

### Experimental section

#### LacCer-BODIPY synthesis and BSA complex preparation

Details of the synthesis process are described in our previous report (16). Briefly, 3.0 µmol β-D-galactopyranosyl -(1->4)-β-D-glucopyranosyl-(1->1’)-D-erythro-sphingosine was dissolved in 200 µL tetrahydrofuran (THF)/H_2_O (3:1 v/v) and mixed with 3.0 µmol BODIPY C_5_ (Thermo Fisher Scientific, Waltham, MA, USA) dissolved in 100 µL of THF/H_2_O (3:1 v/v). 12.3 µmol 4-(4,6-Dimethoxy-1,3,5-triazin-2-yl)-4-methylmorpholinium chloride (DMT-MM; Tokyo Kasei Inc. Tokyo, Japan) was added, and the mixture was stirred for 2 h at 40°C. The product was subsequently purified using preparative thin-layer chromatography. Dried LacCer-BODIPY was dissolved in 200 µL of ethanol and added dropwise to 4 mL of Dulbecco’s phosphate-buffered saline (D-PBS) containing fatty acid-free bovine serum albumin (BSA; Sigma-Aldrich Corp. St. Louis, MO, USA) while vortexing. The complex was dialyzed overnight at 4°C against 500 mL of D-PBS and ultra-centrifuged (100,000 x g) at 4°C for 20 min. Next, the supernatant (LacCer-BODIPY-BSA complex) was collected and stored at −20°C in 1.5 mL tubes.

#### Cell culture

CHO-K1 cells were cultured in Ham’s F-12 (Wako, Tokyo, Japan) containing 10% fetal bovine serum (FBS; Thermo Fisher Scientific) and 1% penicillin-streptomycin solution × 100 (Wako) in culture dishes (Greiner, Kremsmilnster, Austria). Cells were passaged at a density of 80% using 0.25 w/v% Trypsin-1 mM EDTA·4Na solution with phenol red (Wako) at two- to three-day intervals.

#### Pulse chase method

CHO-K1 cells were cultured in 35 mm glass-bottom dishes (IWAKI, Tokyo, Japan) to 60% confluency. The LacCer-BODIPY-BSA complex was diluted with cell culture medium to prepare a 2.5 µM solution. After washing the cells once with saline, 2.5 µM LacCer-BODIPY-BSA complex was added dropwise and incubated at 23°C for 1 min. Next, 2 mL of medium was added after washing twice with saline.

#### Live cell imaging

Lysosomes were stained with LysoTracker® Red DND-99 (Thermo Fisher Scientific) for 5 min. Early endosomes were stained with CellLight® Early Endosomes-RFP (Rab5a-RFP) and BacMam 2.0 overnight. Golgi stacks were stained with CellLight® Golgi-RFP (N-acetylgalactosaminyl-transferase 2 (GalNAcT2-RFP), BacMam 2.0, or CellLight® Golgi-GFP (GalNAcT2-GFP), and BacMam 2.0 (Thermo Fisher Scientific) overnight, respectively. The plasmid of human syntaxin 6-mCherry (pRP[Exp]-Neo-CMV>hSTX6[NM_005819.5](ns):mCherry) (VectorBuilder, Chicago, IL, USA) was constructed and introduced into cells using Lipofectamine® 2000 (Thermo Fisher Scientific) to stain TGN. LacCer-BODIPY was pulsed into cells and chased using time-lapse imaging with a Nikon A1R (Nikon, Tokyo, Japan) at 37°C in a 5% CO_2_ atmosphere.

#### Confocal microscopy

The confocal microscope OLYMPUS FV1000-D (OLYMPUS, Tokyo, Japan) equipped with a temperature control stage (37°C) and UPLSAPO x60/N.A. 1.20 water immersion objective was used. Excitations were set to 488 nm, and emissions were collected at 519 nm. The confocal microscope Nikon A1R (Nikon, Tokyo, Japan) equipped with a temperature control stage (37°C), Plan Apo VC 60x A WI/N.A. 1.20 water immersion objective was also used for time-lapse imaging. Excitations were set to 488 nm and 561 nm, and emissions were collected at 525-575 nm and 595-645 nm.

## Results

### Fluorescently labeled LacCer-BODIPY glycolipid is recycled

In vivo, the homeostasis of glycolipids includes de novo synthesis and multiple recycling pathways. However, the details of these processes are not yet completely understood. Therefore, we examined the uptake of labeled glycolipids in living cells, their incorporation into the cell membrane, and their subsequent trafficking within the cells. For this purpose, we developed a live-cell imaging method to visualize glycolipid recycling. Conventional methods of intracellular trafficking analysis using LacCer-BODIPY (Fig. 1A) apply temperature control to accumulate the probe in the cell and facilitate microscopic observation and mass spectrometry (MS) analysis. However, tracking probe localization changes in the cell has been difficult. Therefore, we tracked glycolipid probe localization by improving the time resolution for 1 min (pulse) and chasing the probe. Under these conditions, the intracellular localization of the probe was observed for a period from 5 to 120 min (Fig. 1B).

After adding the probe, the cells were incubated for 5, 15, 30, 60, 100, and 120 min and observed under a microscope. Localization on the cell membrane was confirmed at 5 min after LacCer-BODIPY incorporation, and vesicle-like fluorescence aggregation was directly observed under the cell membrane after 15 min. The fluorescent probe was observed to be localized near the nucleus in samples incubated for 30 to 100 min.

Furthermore, localized fluorescent molecules could not be observed on the cell membrane after 100 min of incubation. However, the fluorescent probe localized to the cell membrane after incubation for 120 min. These results indicate that the glycolipid probe recycles from the cell membrane to intracellular organelles and back to the cell surface once every two hours.

### LacCer-BODIPY transport depends on its concentration

We previously showed that LacCer-BODIPY is intracellularly metabolized and undergoes glycosylation and degradation (15). In recent years, we have analyzed structural changes in the probe using liquid chromatography-mass spectrometry (LC-MS) by adding LacCer-BODIPY to CHO-K1 cells for 1 min (pulse) and chasing for 10 min (16). This report showed that the probe addition conditions caused differences in glycosylation.

Therefore, we examined the localization of the probe using lysosomal markers following a 30-min chase. Only 5.6% of the probes localized to lysosomes following a 1-min pulse with LacCer-BODIPY and a chase for 30 min. On the other hand, 28.1% of the probes localized to lysosomes following a 10-min pulse and a 30-min chase (Fig. 2).

**Figure 2.**
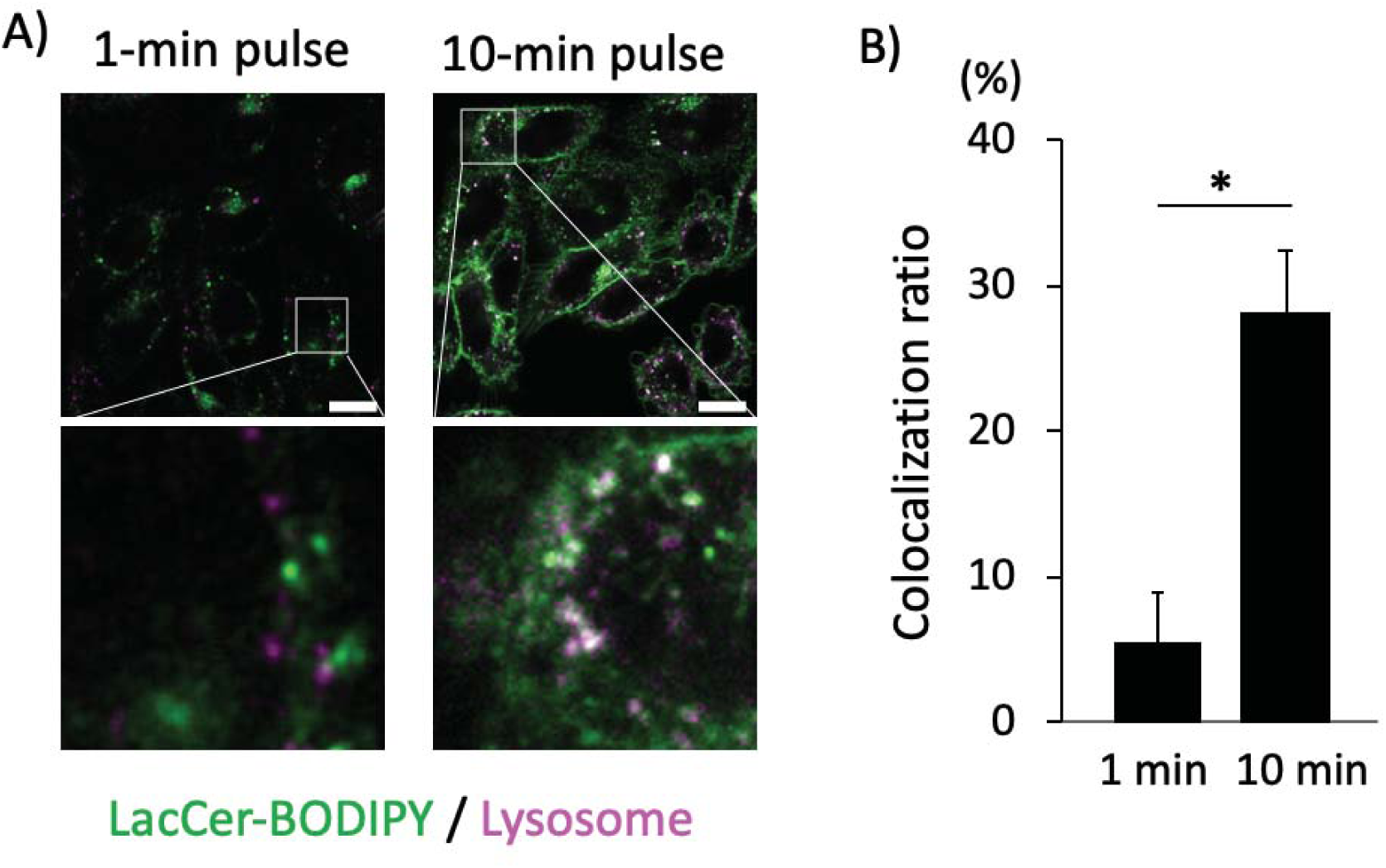
Localization analysis of LacCer-BODIPY with lysosomal markers. A) Localized observation of LacCer-BODIPY to lysosomes following pulse labeling for 1 min (left) and 10 min (right) and chase for 30 min. The lower panels show enlargement of the area within the white-box of the upper panels. Magenta; Lysosome, Green; LacCer-BODIPY Scale bar; 10 µm. B) Co-localization ratio of LacCer-BODIPY to lysosomes. Three cells were randomly selected from the images in this figure, and the ratios of LacCer-BODIPY localized to lysosomes were calculated. Statistical significance was determined using a two-tailed paired t-test. (\**p*<0.003)

This experiment revealed that probe localization is dose-dependent, and this was previously shown using LC-MS analysis. Therefore, these results combined with those presented in Figure 1B suggest that a 1-min pulse with this probe could help visualize the recycling pathway without the lipids being targeted to the degradation pathway.

### Development of a time-lapse imaging method to study LacCer-BODIPY recycling

Time-lapse imaging was performed to examine LacCer-BODIPY recycling in more detail. Conventional observation techniques do not allow continuous observation of the trafficking process because of the fading of BODIPY fluorescence. We used a high-sensitivity photomultiplier tube (PMT), which uses gallium arsenide phosphide (GaAsP) as the photosensitive compound, to suppress fluorescence fading. A photocathode device was the detector in the confocal laser scanning microscope, and the excitation laser power was set as low as possible. We continuously observed glycolipid recycling in one cell by fixing the observation area. Immediately after pulse labeling the cells with LacCer-BODIPY, imaging began at 15 min intervals.

Similar to the results in Figure 1B, it was confirmed that the probe aggregated in the nuclear periphery after 30 min. Thereafter, the probe could not be observed on the cell membrane until 90 min, but it was identified on the cell membrane at 120 min (Fig. 3). The original movie can be found in the supplemental data (Supplemental movie S1).

**Figure 3.**
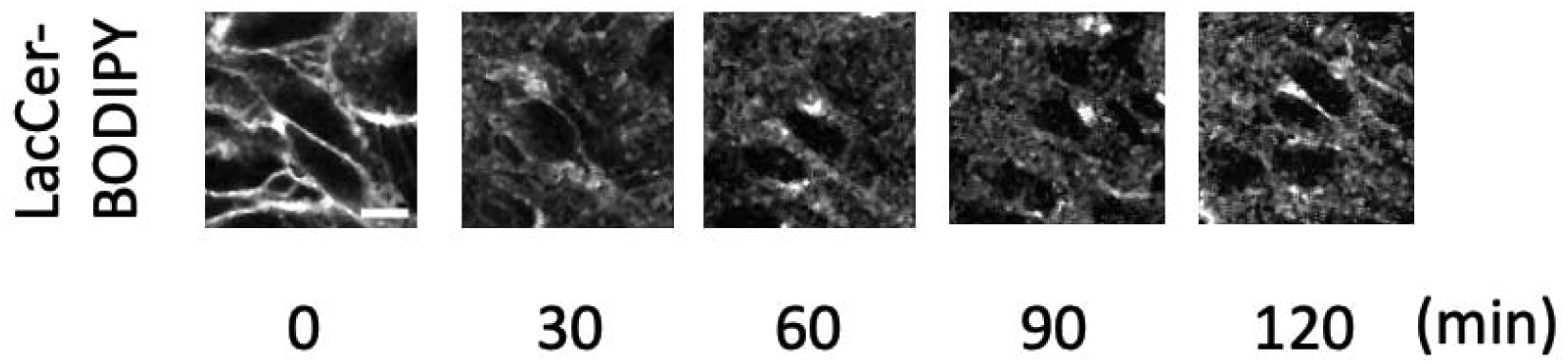
Time-lapse imaging of LacCer-BODIPY using 1-min pulse labeling. 2.5 µM of the probe was taken up into the CHO-K1 cells by pulse labeling for 1 min. The images were taken with a confocal microscope equipped with a GaAsP detector 5 min after pulse labeling. The images were taken every 15 min until 135 min after probe introduction. The images are listed every 30 min. The movie is posted as supplemental data. Scale bar; 10 µm

### LacCer-BODIPY is transported to the TGN via early endosomes

Next, the localization to the early endosome, which is important for endocytosis, was analyzed using time-lapse imaging. It was confirmed that Rab5a-RFP, which is known to localize to early endosomes, and LacCer-BODIPY co-localized after a 30-min chase (Fig. 4A). Furthermore, the same method expressed GalNAcT2-RFP, which is known to be localized in Golgi stacks, in cells. Moreover, its localization with LacCer-BODIPY was examined, but their co-localization was not observed after 60 min and 90 min (Fig. 4B).

**Figure 4.**
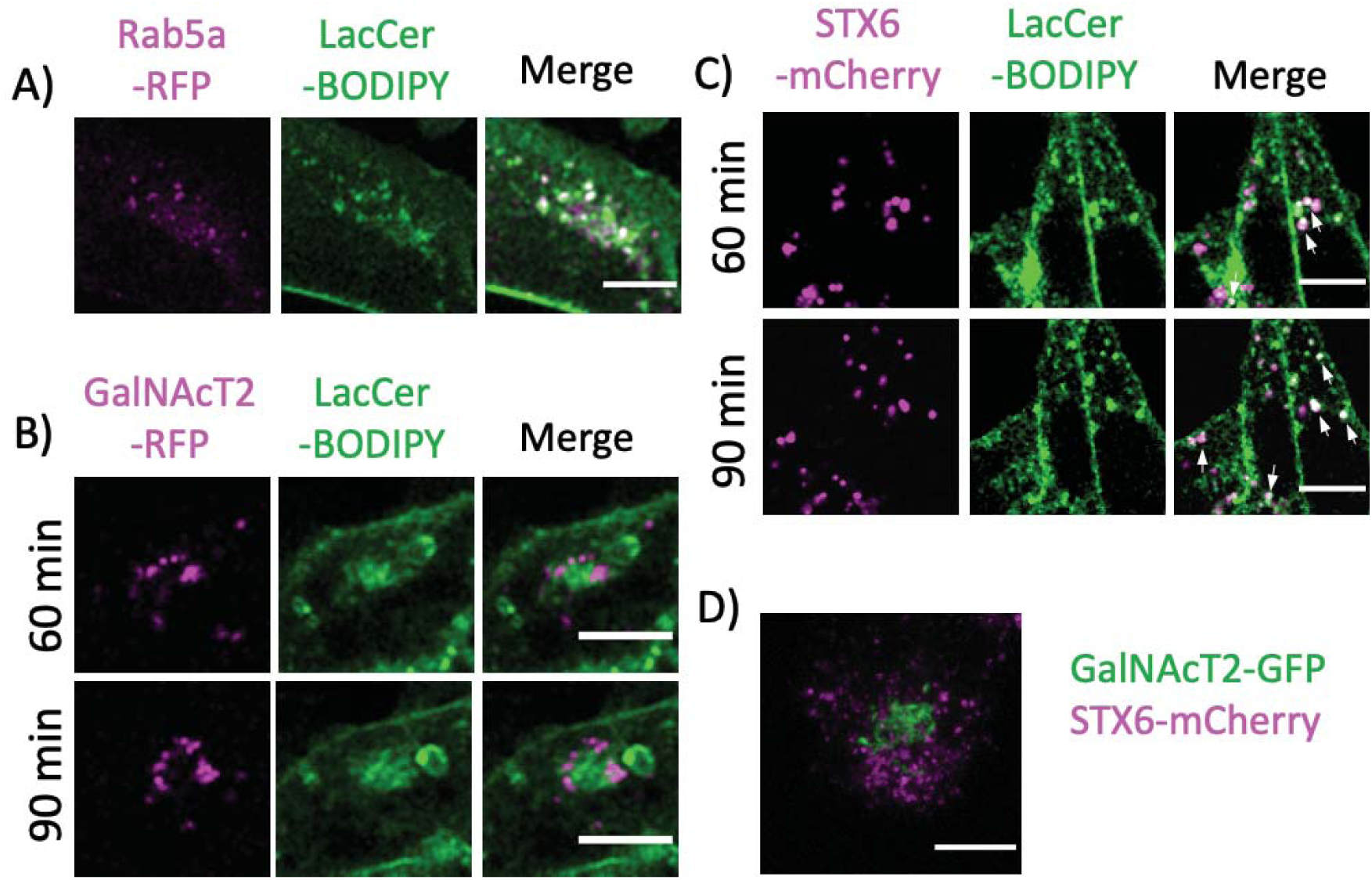
Co-localization of organelle markers and LacCer-BODIPY. A) Localized observation of LacCer-BODIPY in the early endosome. CHO-K1 cells expressing the Rab5a-RFP were pulse-labeled with LacCer-BODIPY and observed after 30 min using confocal microscopy. Magenta; Rab5a-RFP, green; LacCer-BODIPY. B, C) Localized observation of LacCer-BODIPY to the Golgi apparatus. GalNAcT2-RFP expressing cells (B) or STX6-mCherry expressing cells (C) were pulse-labeled with LacCer-BODIPY, and time-lapse observations were performed using confocal microscopy. Magenta; GalNAcT2-RFP (B), STX6-mCherry (C), Green; LacCer-BODIPY. The white arrow indicates co-localization where the pixel color is white. D) Staining of Golgi apparatus with STX6-mCherry (magenta) and GalNAcT2-GFP (green). The view was obtained by performing a standard deviation projection of a Z-stack in ImageJ. Scale bar; 10 µm.

Therefore, we expressed human syntaxin 6, which is known to localize to TGN, fused to mCherry (STX6-mCherry), and examined its co-localization with LacCer-BODIPY. After 60-min and 90-min chases, co-localization of STX6-mCherry and LacCer-BODIPY was observed (Fig. 4C). These results indicated that LacCer-BODIPY was recycled from the early endosome to the plasma membrane via TGN. Furthermore, we also observed that GalNAcT2-GFP and STX6-mCherry did not co-localize (Fig. 4D).

## Discussion

We analyzed the glycolipid transport pathway in more detail using LacCer-BODIPY, which is known as an analysis probe of the glycolipid transport pathway. We successfully visualized the details of glycolipid transport after being taken up from the cell membrane, and this revealed their recycling (Fig. 1B, 3).

Various transport analyses using this probe have been studied by other researchers. In the study of lipid accumulation disease using *Drosophila*, glycolipid uptake by neurons and analysis of their distribution in cells using LacCer-BODIPY have been reported. This study suggested that cholesterol depletion inhibits the transport of glycolipids to the Golgi and ER and promotes their degradation. However, the exact transport pathway in mammals has not been clarified since LacCer is not present in *Drosophila* (17).

Other studies have used mammalian cells. Hoekstra *et al*. observed that fluorescently labeled glycolipids incorporated into BHK-21 cells using liposomes were endocytosed and returned to the cell membrane (18). Pagano *et al*. used BSA to remove residual probes from cell membranes and elucidated the probe transport pathway from early endosomes. However, it was not possible to subdivide the recycling route (12). Here, we first improved the temporal resolution by controlling the amount of probe taken up using a pulse chase (Fig. 1B). It was observed that probe recycling between the cell membrane and the organelle occurred in 2 h (Fig. 1B, 3). In addition, we found that the transport pathway occurs via TGN only in the Golgi apparatus (Fig. 4B-D).

A pathway for the intracellular transport and metabolism of externally taken up glycolipids, involving degradation via lysosomes, has been confirmed in studies using radioisotope-labeled glycolipids (19). Observation of metabolism by glycolipid uptake for 0.5 to 4 h confirmed that most of the introduced glycolipids were transported to lysosomes for degradation (20, 21). Here, we examined glycolipid uptake in a shorter time (1 min). There was less transport of probes to lysosomes with this method. This is the result of a very small amount of tracking compared to previous uptake methods (Fig. 2A left). This is inferred from the fact that, in the previously reported LC-MS study, lysosome-mediated sphingomyelin synthesized by lysosomes was detected by a 10-min pulse, but it was hardly detected by a 1-min pulse (16).

BODIPY is a fluorescent probe that is relatively easy to photobleach. Therefore, it was necessary to reduce the excitation laser power as much as possible and improve the sensitivity of the detector to obtain long-term time-lapse imaging (Fig. 3). We often use a multi-alkali photocathode for the PMT of confocal microscopes. Here, we used the high-sensitivity PMT, GaAsP, which is about ten times more sensitive than a standard PMT (22). This improvement could help detect a relatively small amount of the probe for a long time without significant photobleaching, and this clarified the recycling pathway. Using a similar concept, we previously imaged fluorescently labeled ligands of Toll-like receptor 2 diluted to physiologically active concentrations and successfully visualized receptor expression-dependent endocytosis (23).

The intracellular transport pathway of LacCer-BODIPY identified here is similar to the glycoprotein and glycolipid transport pathways previously investigated using the B subunit of cholera or Shiga toxin (24, 25). These toxins require STX6 for transport and are known to localize to early endosomes before TGN. We demonstrated that LacCer-BODIPY is also transported by a similar pathway. In addition, we were the first to successfully visualize that transport from early endosomes is transferred to the TGN without passing through Golgi stacks.

In conclusion, we visualized the recycling pathway of exogenous glycolipids from early endosomes to TGN by introducing LacCer-BODIPY into cells with short pulses (Fig. 5A). In the previously reported studies, since the glycolipids were taken up in a longer time than the 1-min pulse shown here, we considered that the excess fluorescently labeled glycolipids were transported to lysosomes (Fig. 5B). We visualized the return of lipids to the plasma membrane using time-lapse imaging with this probe. The pathway shown in Figure 5A is likely to rapidly and reasonably return lipids to the plasma membrane to maintain cell membrane homeostasis.

**Figure 5.**
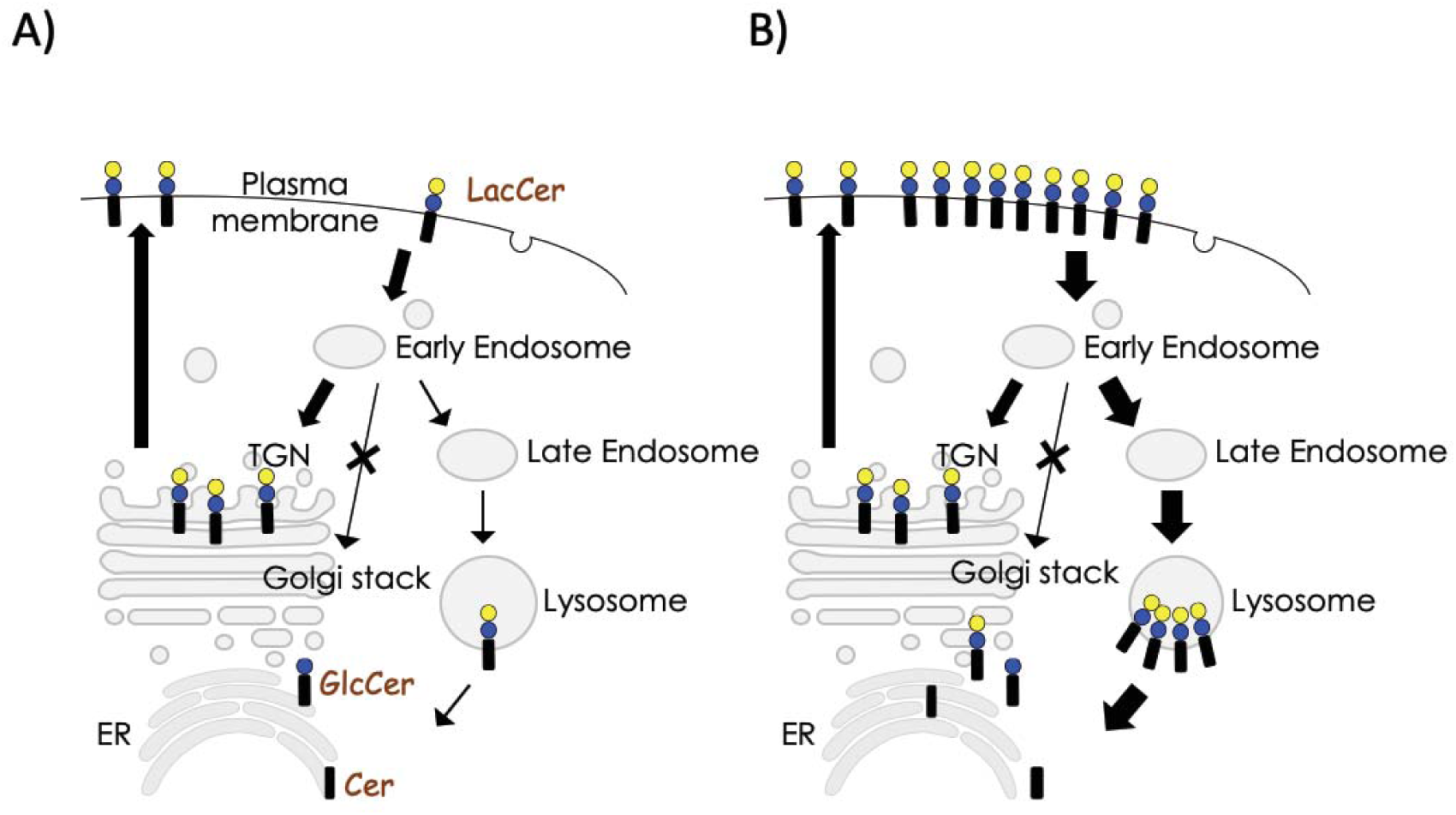
Diagram of the proposed transport pathway for LacCer on plasma membrane. A) Up to a certain amount of LacCer on plasma membrane is transported to early endosomes before returning to the plasma membrane via TGN. B) When large amounts of LacCer are present on plasma membrane, the excess LacCer is transported to lysosome before being degraded or recycled. GlcCer; glucosyl ceramide, Cer; ceramide.

## Abbreviation list

LacCer: lactosyl ceramide,
TGN: trans-Golgi network,
BSA: bovine serum albumin,
THF: tetrahydrofuran,
DMT-MM: 4-(4,6-Dimethoxy-1,3,5-triazin-2-yl)-4-methylmorpholinium chloride,
D-PBS: Dulbecco’s phosphate-buffered saline,
FBS: fetal bovine serum,
GalNAcT2: N-acetylgalactosaminyl-transferase 2,
STX6: syntaxin 6,
MS: mass spectrometry,
LC-MS: liquid chromatography-mass spectrometry,
PMT: photomultiplier tube,
GaAsP: Gallium arsenide phosphide,
ER: endoplasmic reticulum,
GlcCer: glucosyl ceramide,
Cer: ceramide

## Acknowledgements

This work was supported by Grants-in-Aid for Scientific Research, KAKENHI (K. K.; JP 23770155, JP18K05356; K. F.; JP16H01885; O. K.; JP15K14399), by the MEXT-Supported Program (JSPS) for the Strategic Research Foundation at Private Universities (K. K.; S1411010). We would like to thank Editage (www.editage.com) for English language editing.

## Supplemental data

Supplemental movie S1. Time-lapse imaging of LacCer-BODIPY intracellular tracking.

The movie of the intracellular transport of LacCer-BODIPY in CHO-K1 cells was taken using a confocal microscope with GaAsP PMT. This movie was taken every 15 min for 135 min after probe introduction and with automatic brightness correction.

